# Discovering Latent Scientific Concepts through Discrete Representation Learning

**DOI:** 10.64898/2026.10.03.756431

**Authors:** Marjan Stoimchev, Monu Verma, Aveline Filliol, Maria Skamagki, Sara Flowers, Jacob Cohen, Zachary Azadian, Nancy Newlin, Mohamed Saeed Abdel-Mottaleb, Saso Dzeroski, Paul B. Romesser, Scott Lowe, Nevenka Dimitrova

## Abstract

Scientific imaging is undergoing a fundamental transformation. Advances in multiplex immunofluorescence, computational pathology, and electron microscopy now enable observation of biological and physical systems at unprecedented spatial resolution. Rather than simply classifying known objects, these data offer the opportunity to discover previously unknown organizational principles. However, scientific image datasets are typically highly curated, information-rich, and limited in size, making reliable representation learning particularly challenging. We show that standard continuous self-supervised adaptation can become unstable in this regime, allowing teacher representations to drift and degrading downstream transfer.

We introduce Kodiak, a discrete representation learning architecture that discovers *latent scientific concepts* through Sinkhorn-balanced codebook assignments. Instead of learning unconstrained continuous embeddings, Kodiak constructs a reusable vocabulary of discrete concepts that provides stable supervision for both masked-patch prediction and cross-view representation learning within a single-stage framework.

Across five scientific imaging domains, including multiplex immunofluorescence for pancreatic cancer pathology and scanning electron microscopy, Kodiak consistently improves representation quality under both continued foundation-model adaptation and training from scratch. Beyond improved classification and transfer, the learned concepts are interpretable, reusable, and produce spatially coherent concept maps that support applications such as cell phenotyping, segmentation-free patch phenotyping, spatial niche discovery, tissue organization, and materials characterization.

We view Kodiak as the representation-learning engine of our foundation model for spatial biology. More broadly, our results suggest that discrete representation learning provides a principled foundation for discovering reusable latent scientific concepts that can accelerate scientific interpretation across imaging domains.

## 1 Introduction

Scientific imaging is entering a new era. Over the past two decades, biology has been transformed by genomics and single-cell technologies, enabling systematic characterization of genes, proteins, and cellular states. We are now witnessing a complementary transformation driven by spatial biology. Rather than viewing cells as isolated entities, modern imaging technologies reveal that cells exist within spatially organized tissue ecosystems, where neighborhood interactions, microenvironmental context, and higher-order organization govern development, immune function, disease progression, and therapeutic response.

Self-supervised vision foundation models [1,2,3] are reshaping life-sciences imaging. In computational pathology, multiplex immunofluorescence, and microscopy, large encoders—and pathology-specific foundation models such as UNI [4]—now underpin cell typing, tissue characterization, and biomarker discovery. The dominant way to specialize such a model to a new assay, stain, or tissue is to *continue* self-supervised pretraining on specialized domain images [1,5].

Unlike the web-scale corpora these recipes were designed for, however, lifesciences and other scientific datasets are highly curated and information-rich but small: pancreatic multiplex immunofluorescence [4], nanomaterials electron microscopy [6], and environmental imaging [7] typically contain only thousands of images. In these settings, simply continuing self-distillation [3,1] often fails: downstream performance can stagnate or even *degrade* during adaptation. We observe that this instability arises from the *continuous target space* used by modern self-supervised methods [3,8], where teacher embeddings drift when data diversity is limited. To address this, we propose replacing continuous regression targets with *balanced discrete codebook assignments* [9,10], which provide stable categorical anchors for representation learning in specialized domains (Fig. 1).

**Fig. 1:**
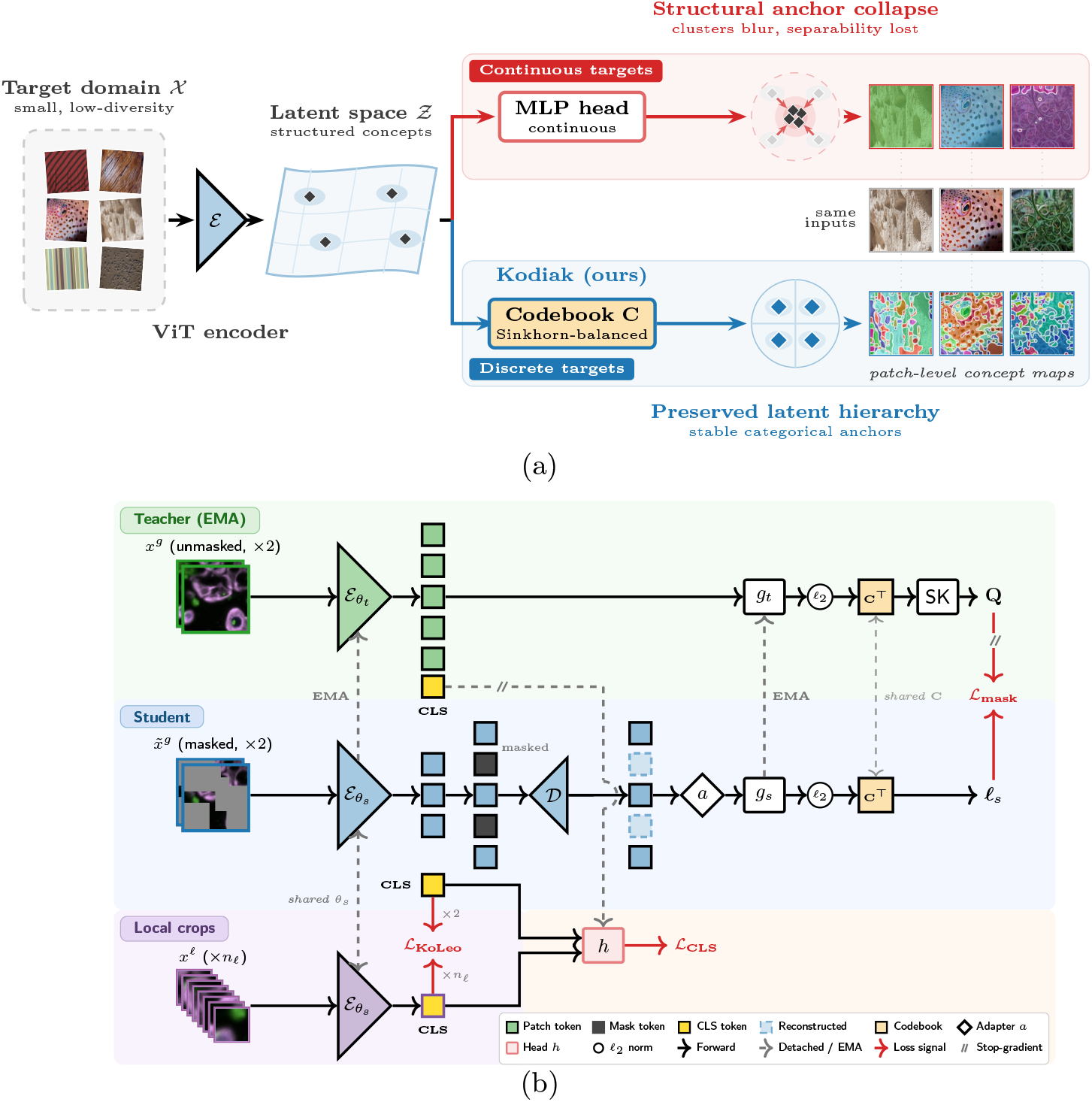
Discrete targets stabilize foundation-model adaptation. **(a)** Under low data diversity the continuous targets of standard self-distillation are weakly constrained, so the representation drifts and cluster structure in latent space *Ƶ* blurs (top); Kodiak’s Sinkhorn-balanced assignments to a shared discrete codebook anchor it on reusable prototypes (bottom). Concept maps at right share the same inputs. **(b)** Kodiak architecture. The EMA teacher maps unmasked global crops to balanced codebook targets **Q**; the student encodes masked crops, reconstructs the patch grid via decoder *D* and adapter *a*, and predicts logits ***l***_*s*_ at masked positions. Head *h* aligns CLS tokens across views; local crops add uniformity signals.

We argue that the next generation of scientific foundation models should not merely optimize predictive performance, but should learn latent scientific concepts: reusable computational representations that capture recurring organization within scientific images and serve as the computational substrate for scientific interpretation and discovery.

### Contributions

Our contributions presented in this paper are as follows:

#### Scientific representation learning

We introduce Kodiak, a discrete representation-learning architecture that discovers reusable latent scientific concepts from complex scientific images.

#### Stable concept discovery

We show that balanced discrete target spaces yield more stable and interpretable concept vocabularies than continuous self-distillation in limited, high-value scientific datasets.

#### Cross-domain scientific generalization

Using multiplex immunofluorescence and scanning electron microscopy, we show that one framework discovers meaningful concepts across distinct scientific domains.

#### Scientific applications

We illustrate how these concepts can support cell phenotyping, segmentation-free patch phenotyping, spatial niche identification, and tumor ecosystem characterization.

## 2 Background and Related Work

### Self-supervised representation learning

Modern self-supervised learning (SSL) has largely evolved along four complementary directions: self-distillation (DINOv3 [11]), masked image reconstruction (MAE [2]), predictive latent modeling (I-JEPA [8]), and masked concept prediction (MOCA [5]). DINOv3 further incorporates Sinkhorn balancing [12] and KoLeo regularization [13] to improve representation quality. Despite their different objectives, these approaches ultimately optimize high-dimensional *continuous* target spaces (except MOCA, which predicts discrete assignments at masked locations). We argue that the stability of this target space becomes a limiting factor in high-value scientific domains with limited training data.

### Discrete representations

Discrete representations have been explored through vector quantization (VQ-VAE [9]) and masked token prediction (BEiT [14]). Unlike these approaches, Kodiak learns a shared Sinkhorn-balanced codebook that simultaneously supervises masked-patch prediction and cross-view alignment within a unified framework, providing a single discrete representation space for both local and global learning.

### Scientific representation learning

Foundation models are increasingly being applied to scientific imaging, including computational pathology through UNI [4]. Unlike these efforts, our objective is not only to adapt foundation models to specialized domains, but to discover stable *latent scientific concepts* that support scientific interpretation across imaging modalities. Kodiak achieves this through balanced discrete representations that remain interpretable, reusable, and transferable across downstream scientific applications.

### Motivation: continuous adaptation can distort the foundation’s geometry

These three lines meet at a practical question: what happens inside a foundation model when it is adapted to a small scientific dataset? To inspect the encoder rather than only its accuracy, we compare each adapted model to the off-the-shelf DINOv3 foundation block by block with linear CKA [15], asking how much of the original layer-wise structure survives. Fig. 2 shows the picture on DTD: continuous-target adaptation does not merely fail to improve the encoder, it visibly reorganizes it, eroding the correspondence with the foundation, whereas discrete balanced targets leave that structure largely intact. This is the failure mode sketched in Fig. 1a—weakly constrained targets let the representation drift until cluster structure blurs—and it is why we treat the *target space*, not the architecture or the recipe, as the variable worth changing. We quantify it in Section 5.1.

**Fig. 2:**
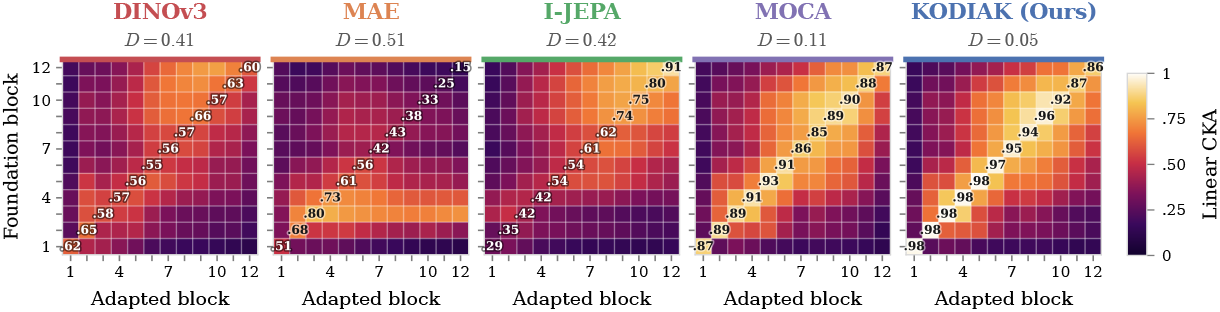
Representation drift under continued pretraining on DTD. Layerwise CKA between each adapted encoder and the off-the-shelf DINOv3 foundation. Cell (*i, j*) is linear Centered Kernel Alignment (CKA) between block *i* of the adapted model and block *j* of the foundation; the drift index 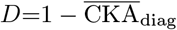 (lower is better) captures same-layer divergence. Kodiak preserves the diagonal; continuous-target methods do not.

## 3 Method

Kodiak retains the core training protocol of DINO [3]—the teacher–student architecture, exponential-moving-average (EMA) teacher updates, multi-crop augmentation, and Vision Transformer backbone. Our modification focuses on a single component: the *target space* used during self-distillation (Fig. 1b). Instead of regressing continuous teacher embeddings, the student predicts Sinkhorn-balanced categorical assignments [12] over a learned visual codebook **C** ∈ ℝ^*K×d*^. This constrains learning to a categorical space of *K* prototypes and empirically mitigates the representational drift described in Section 1. By keeping the architecture and recipe otherwise unchanged, Kodiak isolates the effect of target-space design.

The total objective combines three terms:

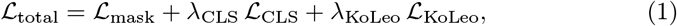

where *λ*_CLS_ and *λ*_KoLeo_ are scalar weights: *L*_mask_ is the core (the student predicts balanced teacher assignments at masked patches), *L*_CLS_ aligns CLS tokens across crops, and *L*_KoLeo_ regularizes for uniformity.

### 3.1 Kodiak Overview

Teacher and student share a ViT-S/16 encoder ε [16] under the DINOv3 configuration, with RoPE positional encoding, register tokens, and LayerScale. The teacher maintains an EMA of student weights and always receives unmasked inputs, while the student follows the MAE sparse forward strategy [2]: only visible tokens enter the encoder given a binary mask **m** ∈ {0, 1}^*N*^. A lightweight transformer decoder *D*reconstructs the full *N*-token grid from visible outputs and learned mask tokens, followed by an adapter *a* (LayerNorm + linear) mapping decoded features back to encoder dimensionality *D*.

#### Patch projection and codebook

MLP projectors *g*_*s*_, *g*_*t*_ map patch features **z** *∈* ℝ^*D*^ into a *d*-dimensional space; the teacher projector is an EMA copy of the student’s. Each projected feature is *l*_2_-normalized, **ū** = *g*(**z**)*/∥g*(**z**)*∥*_2_, before computing cosine logits ***l*** = **ū C**^*⊤*^ *∈* ℝ^*K*^ against the shared codebook **C***∈* ℝ^*K×d*^. The codebook is learned jointly and shared between teacher and student, so both operate in the same prototype space [9].

#### CLS projection head

A separate MLP head *h* : ℝ^*D*^ *→* ℝ^*K*^ projects CLS tokens into a *K*-dimensional categorical space for cross-view alignment (Section 3.3), without routing through **C**. This lets patch- and image-level objectives share categorical dimensionality while specializing their prototype spaces: **C** captures local patch concepts, *h* captures global semantics [3].

### 3.2 Masked Codebook Distillation

The teacher encodes an unmasked global crop and produces patch logits ***l***_*t*_ *∈* ℝ^*N×K*^ against the codebook, which are centered with an EMA offset **c**_p_, temperature-scaled, and converted by Sinkhorn–Knopp [10,12] into balanced soft assignments **Q** = Sinkhorn((***l***_*t*_*−* **c**_p_)*/τ*_*t*_). The Sinkhorn operator enforces an approximate equipartition over prototypes within each batch, preventing code collapse under limited data diversity [10]. The student processes the masked view through the sparse encoder, decoder, and adapter to produce logits ***l***_*s*_ *∈* ℝ^*N×K*^, and the loss is cross-entropy at masked positions *ℐ*_mask_ = *{i* : *m*_*i*_ = 1*}*:

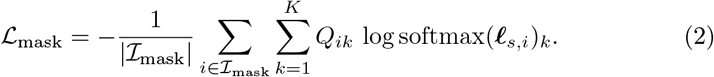

Predicting a *K*-way categorical distribution rather than a continuous teacher vector introduces an explicit information bottleneck that couples masked modeling with balanced clustering, acting as an implicit regularizer against the representational drift analyzed in Section 5.1.

### 3.3 Cross-View CLS Alignment

Following DINO’s multi-crop protocol [3], the teacher processes two unmasked global crops and the student processes masked global and *n*_*l*_ local crops. The shared head *h* projects each CLS token into ℝ^*K*^; teacher targets are centered with EMA offset **c**_c_ and sharpened at temperature *τ*_*t*_, while student predictions use a softer temperature *τ*_*s*_ *> τ*_*t*_, giving 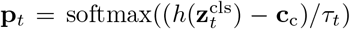 and 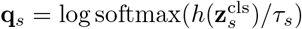. The loss averages cross-entropy over all valid cross-view pairs *P* (global*↔*global and local*→*global, excluding same-crop matches): 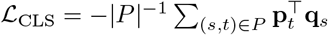. As ablations show (Section 5.7), this regularizer is beneficial when training from scratch but becomes less important when adapting a foundation model that already encodes strong view invariances.

### 3.4 KoLeo Regularization

Following DINOv2/v3 [1], we apply the KoLeo differential-entropy estimator [13] as a uniformity regularizer on *l*_2_-normalized CLS embeddings, extended to both global and local crops: 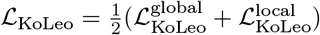, where each per-crop term 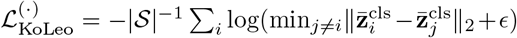 is the standard nearest-neighbor entropy estimate over the corresponding crop set *S*. Separating crop types prevents cross-scale nearest-neighbor pairs from dominating the estimate, providing a complementary safeguard against collapse in continuous embedding space.

## 4 Experimental Setup

We evaluate Kodiak guided by three questions: **(Q1)** does continued selfsupervised pretraining consistently improve foundation models on specialized domains, or can it hinder transfer? **(Q2)** can replacing continuous regression targets with discrete, balanced codebook assignments mitigate such failures? **(Q3)** which components matter, and does their impact vary across training regimes? To disentangle objective design from initialization, we evaluate all methods under three settings:

▪ **Foundation baseline**. The off-the-shelf DINOv3 ViT-S/16 encoder pretrained on LVD-142M [1], evaluated directly—the ceiling adaptation must exceed to be justified.
† **From-scratch pretraining**. Target-domain data only, from random initialization— a stress test of the raw learning signal each objective extracts.
‡ **Continued pretraining**. The same DINOv3 weights, continuing SSL on the target domain—the practical adaptation scenario.

Comparing ^*°*^ against ^‡^ reveals whether adaptation helps or harms; comparing ^†^ against ^‡^ quantifies how much performance comes from foundation initialization vs. the objective itself.

### Datasets

We evaluate on five domains spanning life-sciences and scientific imaging plus controlled natural-image diagnostics (Table 1): an internal multiplex-immunofluorescence pancreatic-cancer dataset (Pancreatic), scanning electron microscopy of nanostructures (SEM [6]), remote-sensing EuroSAT [7], and two low-data natural-image benchmarks, DTD textures [17] and Oxford-Pets [18]. Pancreatic is an extreme low-data regime with high inter-class similarity among fine-grained cell types; DTD and Oxford-Pets are deliberately fragile low-diversity diagnostics where continuous targets are most likely to drift; EuroSAT is a higher-data, lower-class-count reference.

**Table 1:**
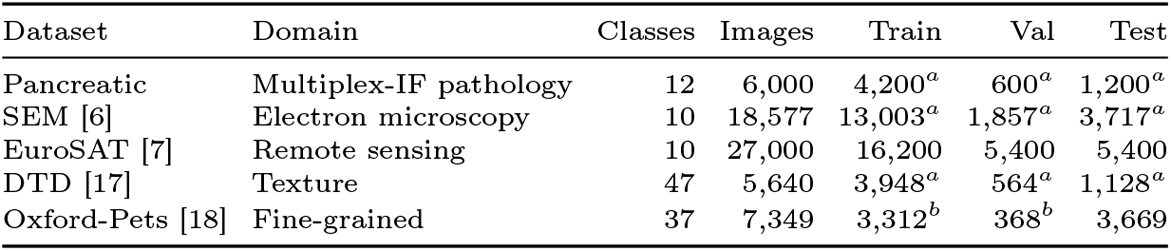
Evaluation datasets. Splits follow official partitions where available; otherwise a fixed-seed 70/10/20 ( ^*a*^) or a 10% validation split carved from official training ( ^*b*^).

### Evaluation protocols

We report four protocols spanning a spectrum of adaptation strength: *k*-NN (*k*=20, *τ* =0.07) and linear probing (LP) [3,1] evaluate frozen-encoder quality; low-shot classification (1–16 labeled examples/class) [5,4] probes label efficiency; and full fine-tuning (FT) [2,14] measures end-to-end performance. This range distinguishes gains from better frozen features from a merely more favorable fine-tuning landscape. To avoid conflating objective quality with FT tuning, we adopt a layer-wise learning-rate-decay FT protocol; Kodiak additionally satisfies FT *>* LP without such tuning (Section 5.6).

### Baselines

We compare against four self-supervised methods spanning pixel-, feature-, and categorical-target objectives: DINOv3 (continuous self-distillation with iBOT-style masked prediction; its checkpoint is also the foundation model), MAE [2] (pixel reconstruction), I-JEPA [8] (latent prediction), and MOCA [5] (codebook assignment without Sinkhorn balancing). All methods share the ViTS/16 backbone and training budget; each keeps its native pretraining recipe, while downstream evaluation is unified. For continued pretraining, all methods start from the same DINOv3 weights. We train with AdamW (batch 128, bf16), two global (256^2^) and eight local (112^2^) crops, and block masking ratio [0.1, 0.5], using *K*=128 codebook entries (*d*=256), chosen as a compromise between representational capacity and prototype utilization; Sec. 5.7 shows that performance is largely insensitive to the exact value of *K*; from-scratch runs use 500 epochs (lr 10^*−*4^) and continued pretraining 100 epochs (lr 10^*−*5^), with EMA-teacher momentum, KoLeo weight 0.1, and CLS weight 1.0.

## 5 Results and Analysis

### 5.1 Continuous Adaptation Leads to Representation Drift

We now put a number on the drift previewed in Section 2. Summarizing each CKA map by 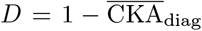 (lower is better) orders the methods cleanly. Continuous-target methods drift strongly (*D≈* 0.41–0.51 for DINOv3, I-JEPA, and MAE), coinciding for DINOv3 and MAE with frozen-feature collapse in Table 2. Kodiak stays anchored at *D≈* 0.05, preserving the block structure of the foundation, while MOCA is intermediate (*D ≈* 0.11). Across methods, larger drift generally correlates with weaker linear-probing accuracy, suggesting that harmful adaptation arises from geometric distortion of the representation rather than optimization issues.

**Table 2:** Main results. *k*-NN, LP, and FT accuracy (%) with ViT-S/16 on five domains (life-sciences and scientific imaging first) under from-scratch (^†^) and continued (^‡^) pretraining. **Best** and second-best among self-supervised methods per regime; gray columns show the frozen foundation (^*°*^) and supervised baseline (^†^). Avg. Rank is over all five domains (lower is better; ranking includes the frozen foundation).

| | | $^\circ$ Fdn. | Sup. † | DINOv3 | MAE | I-JEPA | MOCA | KODIAK (Ours) |
| --- | --- | --- | --- | --- | --- | --- | --- | --- |
| † Pancreatic | $k$-NN | 91.33 | — | **98.66** | 88.04 | 74.26 | 87.70 | <u>97.85</u> |
|  | LP | 95.83 | — | **98.92** | 83.87 | 80.98 | 82.80 | <u>97.28</u> |
|  | FT | 98.52 | 84.74 | **99.53** | 92.34 | 85.82 | 94.89 | <u>97.58</u> |
| † SEM | $k$-NN | 96.84 | — | **94.90** | 83.25 | 70.14 | 80.45 | <u>91.32</u> |
|  | LP | 92.75 | — | <u>76.61</u> | 75.37 | 74.96 | 75.16 | **77.84** |
|  | FT | 95.87 | 70.07 | 88.09 | 87.91 | 86.68 | **90.04** | <u>89.21</u> |
| † EuroSAT | $k$-NN | 94.95 | — | **98.59** | 86.02 | 86.35 | 94.63 | <u>96.94</u> |
|  | LP | 93.06 | — | 92.76 | 72.01 | 82.62 | <u>93.86</u> | **94.56** |
|  | FT | 97.32 | 82.87 | 96.50 | 96.44 | 94.97 | **98.37** | <u>97.57</u> |
| † DTD | $k$-NN | 79.86 | — | 13.05 | 2.69 | <u>25.16</u> | 25.13 | **46.03** |
|  | LP | 79.42 | — | <u>36.49</u> | 1.95 | 23.04 | 26.06 | **47.53** |
|  | FT | 71.12 | 8.47 | 10.45 | 12.42 | 24.45 | <u>34.93</u> | **49.69** |
| † Ox-Pets | $k$-NN | 92.93 | — | <u>7.95</u> | 2.59 | 7.30 | 6.36 | **10.12** |
|  | LP | 93.23 | — | 9.54 | 2.72 | 8.67 | <u>10.78</u> | **12.60** |
|  | FT | 86.22 | 8.62 | 10.84 | 11.17 | 9.37 | <u>15.65</u> | **21.80** |
| Avg. Rank↓ | $k$-NN | 1.80 | — | 2.40 | 5.20 | 4.80 | 4.60 | **2.20** |
|  | LP | 1.80 | — | <u>3.00</u> | 5.20 | 5.40 | 3.80 | **1.80** |
|  | FT | 1.60 | 7.00 | 4.00 | 4.80 | 5.60 | <u>2.60</u> | **2.40** |
| ‡ Pancreatic | $k$-NN | 91.33 | — | 89.65 | 93.08 | **94.62** | <u>93.15</u> | 86.83 |
|  | LP | 95.83 | — | 93.88 | 90.66 | 86.16 | <u>94.09</u> | **94.42** |
|  | FT | 98.52 | — | 96.75 | 96.37 | 97.31 | **98.52** | <u>98.25</u> |
| ‡ SEM | $k$-NN | 96.84 | — | 95.12 | 76.62 | <u>95.67</u> | **96.83** | 95.04 |
|  | LP | 92.75 | — | 89.64 | 76.50 | 90.51 | <u>92.52</u> | **92.81** |
|  | FT | 95.87 | — | 95.28 | 94.05 | 95.32 | <u>95.82</u> | **96.10** |
| ‡ EuroSAT | $k$-NN | 94.95 | — | <u>97.52</u> | 80.61 | 96.26 | 97.15 | **98.59** |
|  | LP | 93.06 | — | 94.33 | 65.25 | 96.45 | <u>97.28</u> | **97.89** |
|  | FT | 97.32 | — | 98.31 | 97.74 | 98.69 | <u>98.77</u> | **98.93** |
| ‡ DTD | $k$-NN | 79.86 | — | 60.41 | 29.14 | **77.68** | <u>77.00</u> | 69.59 |
|  | LP | 79.42 | — | 42.84 | 36.11 | <u>76.84</u> | **76.88** | 75.20 |
|  | FT | 71.12 | — | 62.44 | 73.39 | 75.99 | <u>77.03</u> | **78.00** |
| ‡ Ox-Pets | $k$-NN | 92.93 | — | 67.90 | 21.27 | 78.17 | **83.75** | <u>80.00</u> |
|  | LP | 93.23 | — | 20.64 | 32.93 | 88.24 | <u>90.58</u> | **91.03** |
|  | FT | 86.22 | — | 80.26 | 88.16 | 88.12 | <u>89.58</u> | **91.61** |
| Avg. Rank↓ | $k$-NN | 2.40 | — | 4.20 | 5.40 | <u>2.80</u> | **2.40** | 3.80 |
|  | LP | 2.00 | — | 4.80 | 5.60 | 4.00 | <u>2.60</u> | **2.00** |
|  | FT | 3.90 | — | 5.20 | 4.80 | 3.60 | <u>2.10</u> | **1.40** |

### 5.2 Kodiak Discovers Stable Latent Scientific Concepts

A key promise of discrete targets for science is that the codebook yields a vocabulary of reusable, inspectable visual concepts. Fig. 3 visualizes patch-level concept assignments under continued pretraining. Kodiak’s prototypes emerge and specialize, producing spatially coherent regions without any post-hoc clustering: texture variations in DTD, object parts in Oxford-Pets, land cover in EuroSAT, nanostructure motifs in SEM, and cell/compartment structure in Pancreatic. Continuous-target baselines (I-JEPA, MAE, DINOv3) lack a native prototype vocabulary and rely on *K*-means over frozen features, collapsing to one or two dominant regions; MOCA produces multi-concept maps but exhibits a dominant prototype, reflecting the absence of Sinkhorn balancing. As a quantitative check, a frozen linear segmentation probe on Oxford-Pet trimaps ranks Kodiak best among continued-pretraining methods (97.5% mIoU, *−* 0.2 pp from the foundation) ahead of I-JEPA (97.1), MOCA (95.6), DINOv3 (93.2), and MAE (84.2), mirroring the classification ranking. These observations align with the CKA analysis (Section 5.1): discrete balanced prototypes preserve local structure during adaptation.

**Fig. 3:**
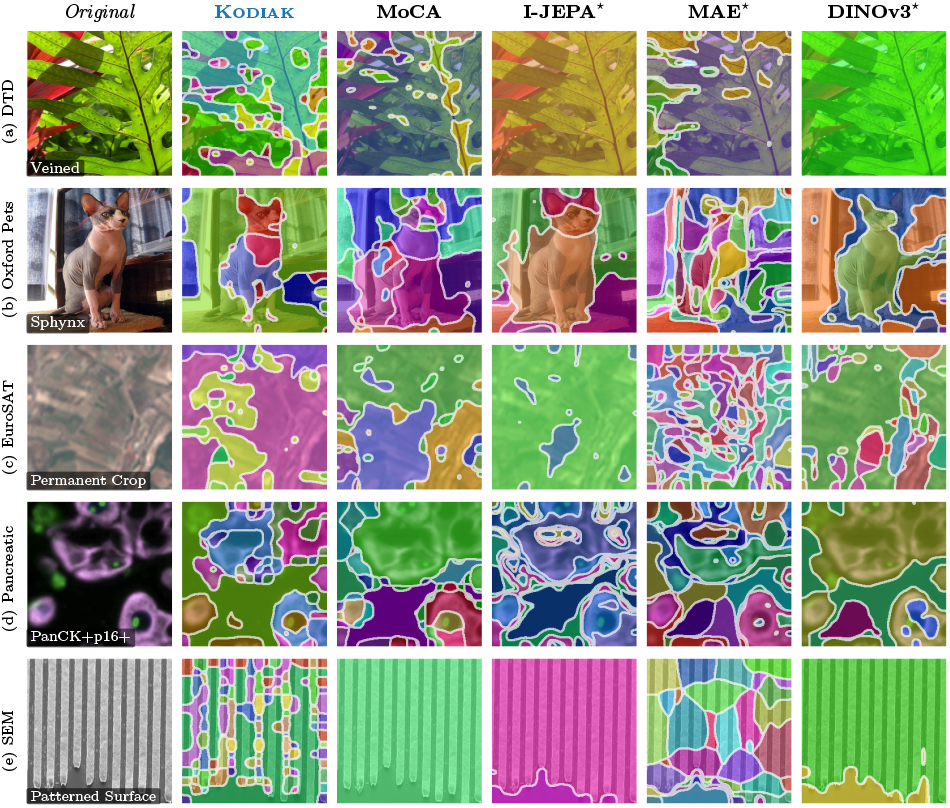
Patch-level concept segmentation. (continued pretraining). Kodiak captures fine-grained local structure while continuous-target baselines collapse to a dominant cluster. ^*⋆*^ I-JEPA, MAE, and DINOv3 use *K*-means (*K*=128) on frozen patch features; Kodiak and MOCA use the learned codebook directly.

### 5.3 The Continuous Geometry Behind the Concepts

Concept maps show how the codebook *partitions* an image; projecting the dense patch features to RGB via a per-image three-component PCA shows the continuous geometry underneath [11]. Fig. 4 shows the outcome across the five domains. Kodiak and MOCA produce spatially coherent maps whose boundaries track semantically meaningful regions—object silhouettes on Oxford-Pets, texture boundaries on DTD, land-cover transitions on EuroSAT, and cell-type regions on Pancreatic. MAE is smooth but semantically undifferentiated, consistent with its pixel-level objective, and I-JEPA is intermediate; adapted DINOv3 keeps coarse spatial coherence but with weaker domain-specific boundaries. The discrete partition and the continuous geometry therefore agree, both indicating that Kodiak preserves local structure during adaptation.

**Fig. 4:**
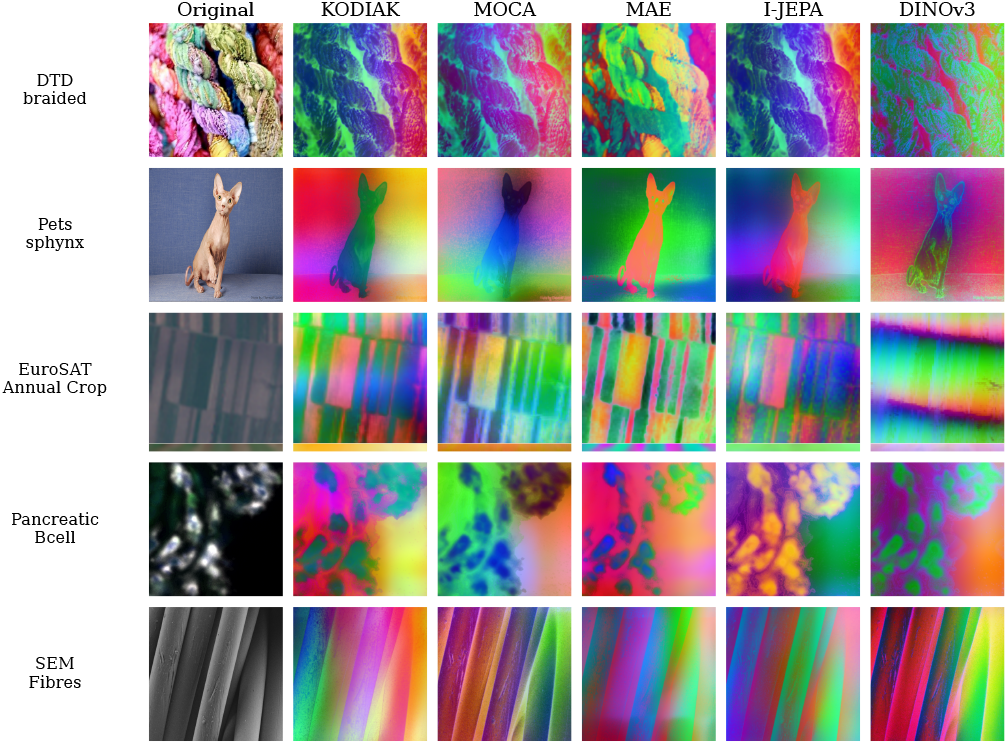
PCA feature visualization. (continued pretraining, ViT-S/16). Per-image PCA over dense patch tokens, mapped to RGB. Kodiak and MOCA give spatially coherent maps with clear semantic boundaries; MAE is smoother and less structured.

### 5.4 Latent Scientific Concepts Improve Scientific Representation Learning

Table 2 reports *k*-NN, LP, and FT accuracy across both regimes and all five domains, asking whether the latent concepts of Section 5.2 translate into quantitative gains.

#### Discrete balanced targets preserve transferable structure

Under continued pretraining, Kodiak achieves the best fine-tuning performance on DTD (78.0%), Oxford-Pets (91.6%), EuroSAT (98.9%), and SEM (96.1%). Where DINOv3 collapses frozen linear separability (93.2 *→* 20.6% LP on Oxford-Pets; 79.4 *→* 42.8% on DTD), Kodiak recovers much of the foundation geometry and exceeds the frozen foundation under fine-tuning. On the biomedical Pancreatic data, where continuous adaptation is less catastrophic, Kodiak remains the best continued-PT method under linear probing (94.4%) and top-2 under LP and FT, indicating stable adaptation rather than domain-specific overfitting.

#### From-scratch training exposes objective quality

From random initialization, Kodiak substantially outperforms all baselines on the low-data diagnostic domains (DTD, Oxford-Pets). On DTD, pixel reconstruction collapses (MAE LP 2.0%, near chance for 47 classes) and feature-level objectives trail far behind (I-JEPA LP 23.0% vs. 47.5%). On SEM, Kodiak attains the highest from-scratch LP among self-supervised methods (77.8%), while DINOv3 remains ahead on Pancreatic. These results indicate that a discrete, balanced target space provides a stronger learning signal from limited scientific data than continuous regression, even without foundation-model priors.

#### Performance gains are broad but regime-dependent

Across all five domains, Kodiak attains the best average rank under LP and FT in both regimes (Table 2). The largest gains occur where data are limited, semantically dense, or visually repetitive—where drift is most harmful—and shrink on higher-data or already well-structured domains, indicating that discrete concepts matter most when datasets are curated, information-rich, and insufficient to constrain continuous targets.

### 5.5 Latent Scientific Concepts Generalize Under Limited Supervision

Annotation in the life sciences is expensive, so we evaluate low-shot transfer (1–16 labeled examples/class) under fine-tuning and linear probing (Fig. 5). Under fine-tuning, Kodiak leads across shot budgets on the scientific and fine-grained domains (Pancreatic, SEM, DTD, Oxford-Pets), with the largest margins on Pancreatic and SEM, and stays competitive on EuroSAT. Under linear probing it leads on Oxford-Pets, EuroSAT, Pancreatic, and SEM at all budgets, while DINOv3 collapses under LP on DTD and Oxford-Pets even at 16 shots—confirming its degradation is frozen-feature drift, not an optimization failure.

**Fig. 5:**
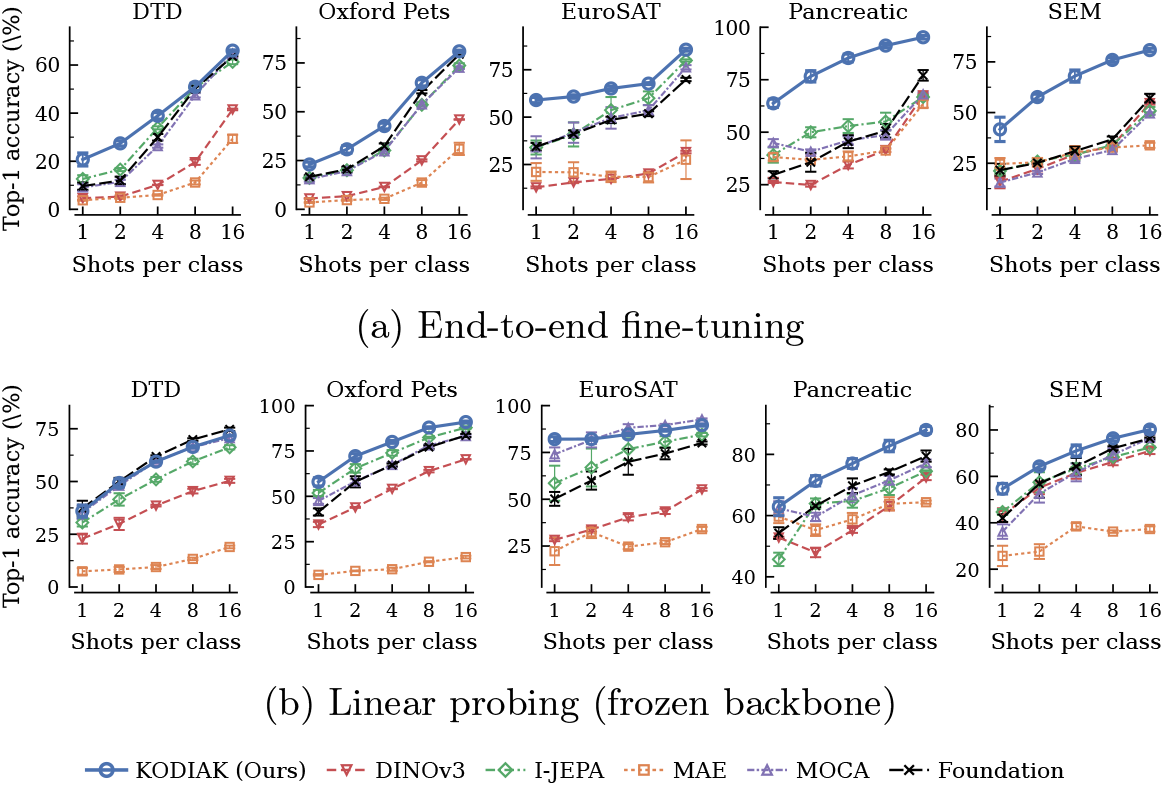
Low-shot classification. (continued pretraining, ^‡^). Top-1 accuracy vs. labeled examples per class (1–16) under (a) end-to-end fine-tuning and (b) linear probing; mean*±* std over 3 label seeds. ^*°*^ *Foundation*: off-the-shelf DINOv3, no adaptation. Kodiak leads across nearly all shot budgets on the scientific and fine-grained domains.

### 5.6 Stable Adaptation, Not Universal Improvement

Continued pretraining does not universally beat the frozen foundation, and we report this transparently by measuring, for every continued method, the change relative to the frozen foundation across all five domains. For frozen protocols the foundation is a strong baseline, and the right goal is *stable* adaptation. Two points stand out. First, on linear probing, where the failure mode is most severe, Kodiak averages only *−* 0.6 pp vs. DINOv3’s *−* 22.6 pp—a ~38 *×* reduction in frozen-feature drift. Second, under fine-tuning—the protocol practitioners actually use—Kodiak achieves the best average gain over the foundation (+2.8 pp), with large improvements on DTD (+6.9) and Oxford-Pets (+5.4). Kodiak thus trades a small frozen-feature cost for the strongest, safest end-to-end adaptation. The same robustness holds under FT tuning: Kodiak satisfies FT *>* LP under the original protocol without layer-wise decay, whereas the foundation requires it, confirming Kodiak’s advantage is not a tuning artifact.

### 5.7 Understanding the Role of Discrete Target Spaces

#### Component importance is regime-dependent

Removing *L*_CLS_ or *L*_KoLeo_ from scratch causes large drops on DTD (*>*24 points; Table 3a), showing these auxiliary signals stabilize learning in the absence of pretrained geometry. Under continued pretraining the trend reverses: the discrete codebook objective captures most gains, and *L*_CLS_ can even be mildly constraining on Oxford-Pets (+1.5 without it). The practical recipe: use the full objective when training from scratch, and consider patch-only codebook distillation when adapting strong foundations.

**Table 3:**
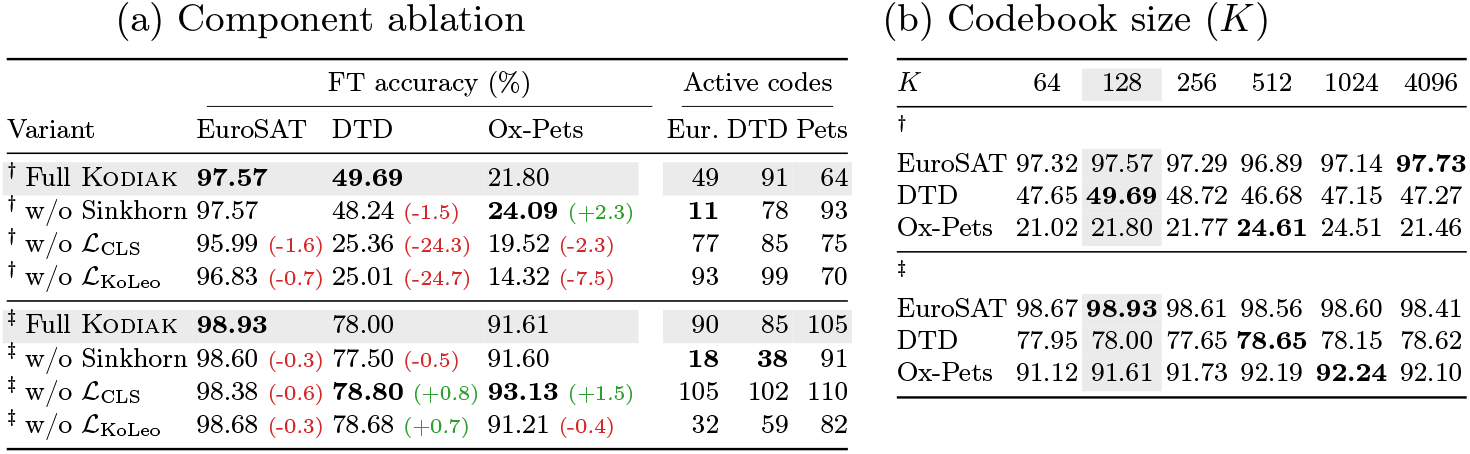
Design analysis. (a) Component ablation: FT accuracy (%) and active codebook entries (of *K*=128) under both regimes; green/red deltas vs. full Kodiak. (b) Codebook size (*K*) sweep. Gray: default (*K*=128). ^†^: from scratch. ^‡^: continued PT.

#### What Sinkhorn balancing does, and what it does not

The “w/o Sinkhorn” variant in Table 3a remains fully discrete: it retains the learned codebook **C** and masked codebook objective, replacing only the balanced assignments **Q** with a plain softmax over the same logits. Its competitive accuracy shows that the primary benefit comes from the *discrete target space* itself rather than from Sinkhorn balancing. Removing Sinkhorn reduces downstream accuracy by at most 0.5 pp under continued pretraining.

Sinkhorn instead plays a different role. It substantially increases *prototype utilization*, encouraging the model to distribute assignments across a larger fraction of the available codebook. On low-diversity domains, removing Sinkhorn reduces the number of active prototypes from 90 to 18 on EuroSAT and from 85 to 38 on DTD, while leaving classification performance largely unchanged. Where collapse occurs, it is pronounced: in the DTD ablation, the top 10% of prototypes account for 96% of all assignments, compared to 55% for the full model, effectively shrinking the learned vocabulary to only a dozen concepts. In contrast, Oxford-Pets exhibits little change, suggesting that datasets with greater intrinsic diversity naturally sustain broader codebook usage.

These results indicate that Sinkhorn is not the primary source of predictive performance. Rather, it promotes a richer and more balanced concept vocabulary, which is important when the objective extends beyond classification to discovering reusable latent scientific concepts.

#### Codebook size

Performance is stable across *K* ∈ [64, 4096] (Table 3b), so *K* is not accuracy-critical; we adopt *K*=128 as the smallest setting still yielding a vocabulary of order 10^2^ active concepts—expressive enough for these domains, small enough to inspect prototype by prototype.

## 6 Discussion

### Why discrete targets enable structure-preserving adaptation

The CKA analysis (Section 5.1) supports our hypothesis mechanistically. Continuous regression targets let mid-to-late layers drift from the foundation, collapsing frozen-feature separability. Discrete codebook prediction introduces a categorical bottleneck that anchors features to a finite set of prototypes, stabilizing adaptation. Sinkhorn balancing plays a narrower role than the ablation accuracies alone would suggest: the categorical bottleneck, not the balancing, is what stabilizes adaptation, while Sinkhorn determines whether the resulting codebook remains a usable vocabulary instead of concentrating onto a few dominant prototypes (Section 5.7).

### Implications for the life sciences

Although this work focuses on robust representation learning, we envision the discovered concepts serving as the computational substrate for future models of cell phenotypes, spatial ecosystems, and tissue organization across imaging modalities.

For practitioners adapting foundation models to scarce, curated assays, Kodiak offers a safe default: it never catastrophically degrades the foundation, leads under the low-shot budgets typical of expert annotation, and exposes a reusable concept vocabulary that supports inspection and discovery, not only prediction.

### Scope and limitations

Results are single-seed, so small gaps between closely matched variants should be read as indicative rather than significant, and the CKA analysis shows that drift co-occurs with degraded frozen transfer rather than establishing causation. We evaluate at ViT-S/16 throughout, which keeps the comparison controlled but leaves the behaviour of discrete targets at larger backbone scales an open question. As domain data grow larger and more diverse, the advantage of discrete targets narrows, since continuous targets become better constrained. We study classification transfer, though the codebook already induces coherent segmentations (Section 5.2), suggesting benefits for dense prediction.

## 7 Conclusions

We introduced Kodiak, a discrete representation learning architecture that discovers reusable latent scientific concepts from complex scientific images. By replacing continuous targets with balanced discrete codebook assignments, Kodiak stabilizes representation learning across scientific domains while improving transfer and concept interpretability.

Our results show that the primary benefit comes from the discrete target space, whereas Sinkhorn balancing promotes a richer and more balanced concept vocabulary. Beyond classification, the learned concepts support scientific applications including cell phenotyping, segmentation-free patch phenotyping, spatial niche discovery, and materials characterization.

We view Kodiak as the representation-learning engine of the *Second Foundation (2ndF)* and believe that learning reusable latent scientific concepts provides a promising foundation for future scientific discovery.

## Acknowledgments

SD and MS were supported by the Slovenian Research and Innovation Agency (ARIS) via the research program P2-0103 and the research projects GC-0001 and GC-0006. AF, SL, PR were supported by the *Mark Foundation for Cancer Research Endeavor Award*.

## Disclosure of Interests

The authors have no competing interests to declare that are relevant to the content of this article.

## Data availability and ethics

Code for Kodiak will be released at https://github.com/marjanstoimchev/KODIAK. Public benchmarks (SEM, EuroSAT, DTD, Oxford-Pets) are available from their original sources. The internal pancreatic multiplex-immunofluorescence dataset was collected under institutional approval and is available from the authors upon reasonable request, subject to data-governance constraints.

## Notes

### Competing Interest Statement

The authors have declared no competing interest.

